# Multiplicative Joint Coding in Preparatory Activity for Reaching Sequence in Macaque Motor Cortex

**DOI:** 10.1101/2023.04.05.535305

**Authors:** Tianwei Wang, Yun Chen, Yiheng Zhang, He Cui

## Abstract

Although motor cortex has been found to be modulated by sensory or cognitive sequences, the linkage between multiple movement elements and sequence-related responses is not yet understood. Here, we recorded neuronal activity from the motor cortex with implanted micro-electrode arrays and single electrodes while monkeys performed a double-reach task that was instructed by simultaneously presented memorized cues. We found that there existed a substantial multiplicative component jointly tuned to impending and subsequent reaches during preparation, then the coding mechanism transferred to an additive manner during execution. Multiplicative joint coding, which also spontaneously emerged in a recurrent neural network trained for double-reach, enriches neural patterns for sequential movement, and might explain the linear readout of elemental movements.

## Introduction

Motor cortex has long been thought to be central in planning and generating movement. A large body of evidence demonstrates a correlation between neuronal activity in motor cortex and a variety of motor variables, such as direction, speed, distance, and trajectory ^1–7^. Beyond the single ballistic movements examined in these studies, multi-step movements, such as sequencing and ordering action, are crucial in daily behavior ^8, 9^. As one of the brain areas conveying highly accurate information about movement timing ^10^ and kinematics ^11^, motor cortex seems to be involved in causal sequencing of multi-step movements ^12^. Sequential information has been reported to be encoded in the population response before movement initiation ^13–15^. In addition, most neurons are reported to show activity related to both target location and serial order ^16, 17^. However, most of these studies instructed the sequence of movement with serial sensory stimuli, which might result in neural activity that differs from internally generated motor sequences^18–20^. In tasks carried out in the absence of serial sensory inputs, neuronal activity related to sequential contexts emerges during preparation, and becomes prominent during execution ^21, 22^. Furthermore, despite differences at the single-neuron level, the neural population preserves a reliable readout of movement direction. That is to say, both individual movement elements and sequential information are simultaneously and robustly encoded in the motor cortex ^21^.

In principle, a continuous action sequence consists of elements spatio-temporally coordinated in a complex manner, rather than a series of independent actions ^23–25^. However, the ‘competitive queuing’ hypothesis suggests that the brain produces sequential movement via a combination of parallel representations of specific actions ^26^. A recent study on double-reach supports this parallel-representation hypothesis, suggesting that motor cortex does not fuse two reaches, but recruits two independent motor processes sequentially ^27^. The resulting concurrence of motor execution and motor planning, however, is insufficient for rejecting the possibility of interaction between movement elements beforehand. It remains unclear if sequential movement is parallel or jointly coded in the preparation period.

To further explore the motor preparation and encoding characteristics of sequential movements in a strict behavioral and neurophysiological context, we recorded neuronal activity from the motor cortex via implanted arrays or single electrodes while monkeys were performing a double-reach that was instructed by simultaneously presented cues that had to be memorized. We found that neuronal activity could be regressed as a multiplication of directional tunings to reaching elements in the preparatory period, and then converted to parallel coding for both movement elements after movement onset, indicating the existence of a gain-like interaction in planning the motor sequence. Neural population dynamics derived from our array-recorded data indicates that a nonlinear interaction is embodied in the spatial structure of initial states. In computational simulations, multiplicative coding for motor sequences spontaneously emerges in a recurrent dynamical network, and benefits reliable linear readouts of movement elements. These results suggest that the motor cortex is profoundly involved in concatenating multiple movement elements into a sequence, and that a gain-like multiplication is a key signature of complex serial behavior.

## Results

### Behavioral task

Three rhesus monkeys (*Macaca mulatta,* male 5-10 kg) performed the memory-guided double-reach task (Fig. 1a). A trial began with a green dot displayed on the center of a touch screen, and the monkey was required to touch it. After 300 ms, in 1/3 of the trials (single-reach, SR), another green dot was presented as a reaching goal for 400 ms (cue period) at one of the six corners of a regular hexagon (i.e., at directions of 0°, 60°, 120°, 180°, 240°, or 300°). After the peripheral cue was extinguished, there was a memory period of 400-800 ms. The monkey was trained to keep its hand on the central green dot until it was turned off (GO signal), and then reach to the previously cued location to obtain a reward. In the remaining trials (double-reach, DR), a green square and a green triangle were presented simultaneously during the cue period. The square was in the same alternative directions as the SR surrounding targets. The triangle was displaced from the square by 120° clockwise (CW, 1/3 of trials) or 120°counterclockwise (CCW, 1/3 of trials). After the memory period without peripheral cues, the monkey was required first to reach to the memorized square location, and then to immediately reach to the memorized triangle location. The monkey was rewarded only if it reached the specified target within a margin of three centimeters, and in the correct order. For a correct trial, the green square would reappear after the 1st reach, and the triangle would appear in purple after the 2nd reach. All 18 conditions (three trial types × six directions) were pseudo-randomly interleaved. Only correct trials were included in the analysis. Event markers are denoted as the GO signal (GO), the 1st/only movement onset (MO), the 1st/only movement end (ME), and the 2nd movement onset (MO2).

**Figure 1.**
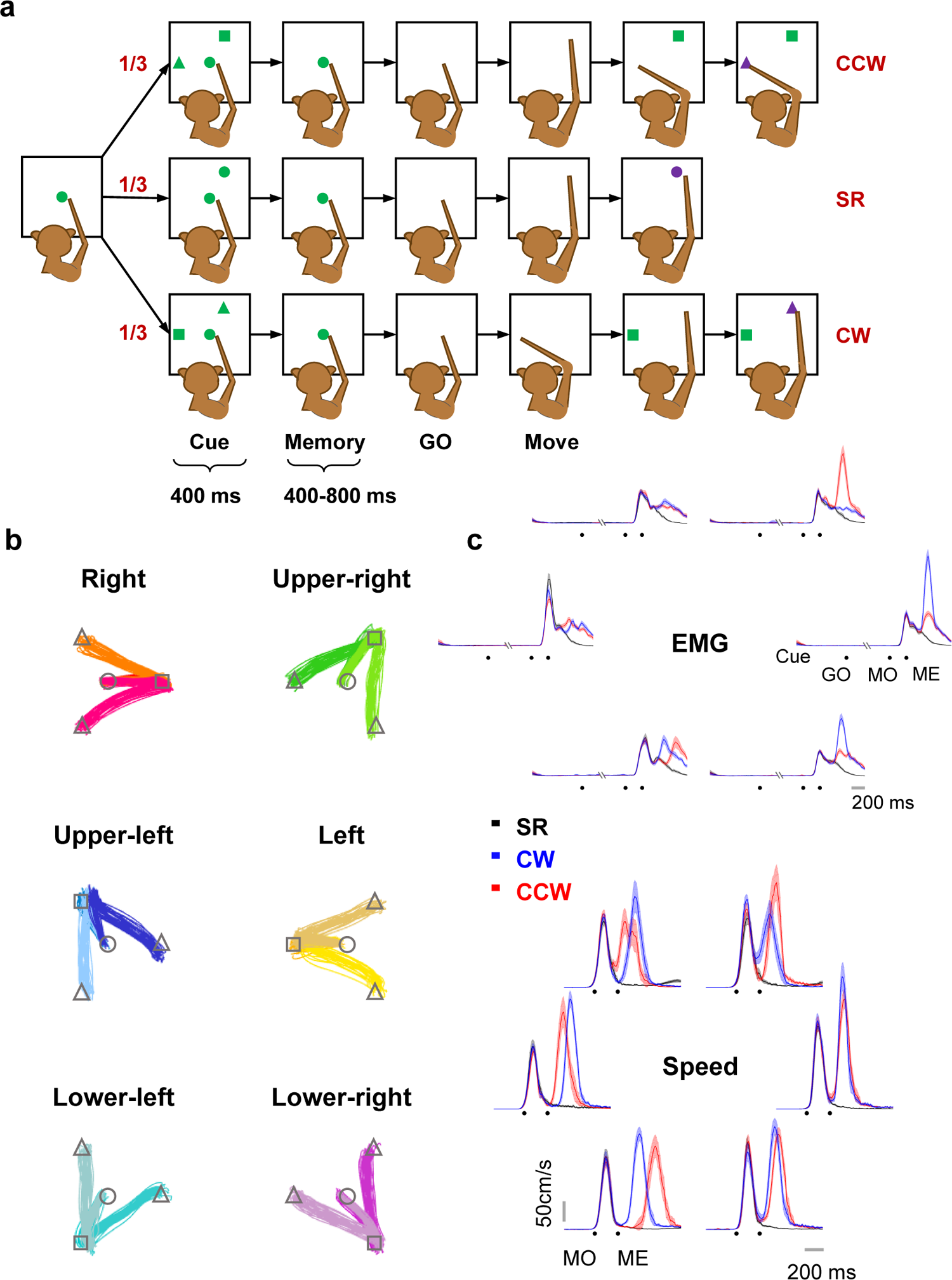
Paradigm and behavior. **a.** Three types of trials were pseudo-randomly interleaved in each session. In single-reach (SR) trials, monkeys had to perform memory-guided center-out reach. In double-reach (DR) trials, two targets (a square and a triangle) were presented simultaneously in cue period, and then extinguished; the monkeys were required to hold the central target for a 400-800 ms memory period until it was turned off (GO signal). Next, monkeys finished reaching both targets in the sequence of the square to the triangle within 700-1200 ms. The triangles were located 120° from the squares in CW or CCW directions. b. Hand trajectories in different conditions are grouped by their 1st/only reach direction from monkey C. Some trajectories are overlapped due to high similarity. No significant difference was found before the end of 1st/only reach (one-way ANOVA, p>0.05). c. Surface electromyography (sEMG) and speed in one typical session. The Pearson correlation coefficient of the speed profile until 1st movement end between double reach and single reach was 0.99±0.006 (mean±sd) and of sEMG of extensor digitorum communis (EDC) was 0.99±0.005 (mean±sd) for monkey C.

Hand trajectories exhibited a stereotype movement pattern in each condition for well-trained monkeys. All 1st reaches started from the center and moved towards the corresponding target in each condition (Fig. 1b). Muscular activities remained constant during the preparatory period across different conditions, excluding the possibility that the monkeys might develop different premature movements (e.g., adjust arm orientation) after cue for different conditions. The Pearson correlation coefficient of speed profiles until ME between DR and SR was 0.99±0.006 (mean±sd), and of surface electromyography (sEMG) of extensor digitorum communis (EDC) was 0.99±0.005 (mean±sd) for monkey C (Fig. 1c). In addition, the dwell time on the 1st target was 194±75 ms (mean±sd) for monkey C, 350±110 ms (mean±sd) for monkey G, and 150±47 ms (mean±sd) for monkey B. The median duration of DR was 586±95 ms (mean±sd) for monkey C, 818±131 ms (mean±sd) for monkey G, and 481±72 ms (mean±sd) for monkey B, averaged across conditions. These results verified the expected transitory dwell on the 1st target in this task, and indicated behavioral consistency between SR and the 1st reach of DR in the same direction, in terms of hand trajectory, speed profile, and sEMG.

### Heterogeneity in neuronal activity indicated mixed selectivity

All electrophysiological recording sites were in the hemisphere contralateral to the hand used during the task. Only one hand was used by monkeys B and G, but for monkey C data were recorded first with single electrodes, and then arrays in the other hemisphere with a switch of hands. We collected 322 well-isolated task-related neurons from single-electrode recordings (224 from monkey B, 98 from monkey C left hemisphere) and 202 units sorted from array recordings (44 from monkey G, 158 from monkey C right hemisphere) in motor cortex (Fig. S1). Among these, we found considerable heterogeneity in firing patterns. Figure 2 illustrates four representative cells. The neuron in Fig. 2a exhibited a two-peak firing pattern in DR, each peak after a movement onset, while it had only one burst in SR. Notably, the direction with the highest firing rate changed remarkably in sequential movements. The neuron in Fig. 2b fired with a constant PD towards the lower left. Surprisingly, even though its directional selectivity was remarkably similar for both SR and DR, the firing rate was significantly higher in DR (according to the 95% confidential interval plotted in shade), indicating that it conveyed information regarding target-movement number. Also, the preparatory activity would diverge with the 2nd reach before GO and MO in neurons, as in Fig. 2c and 2d.

**Figure 2.**
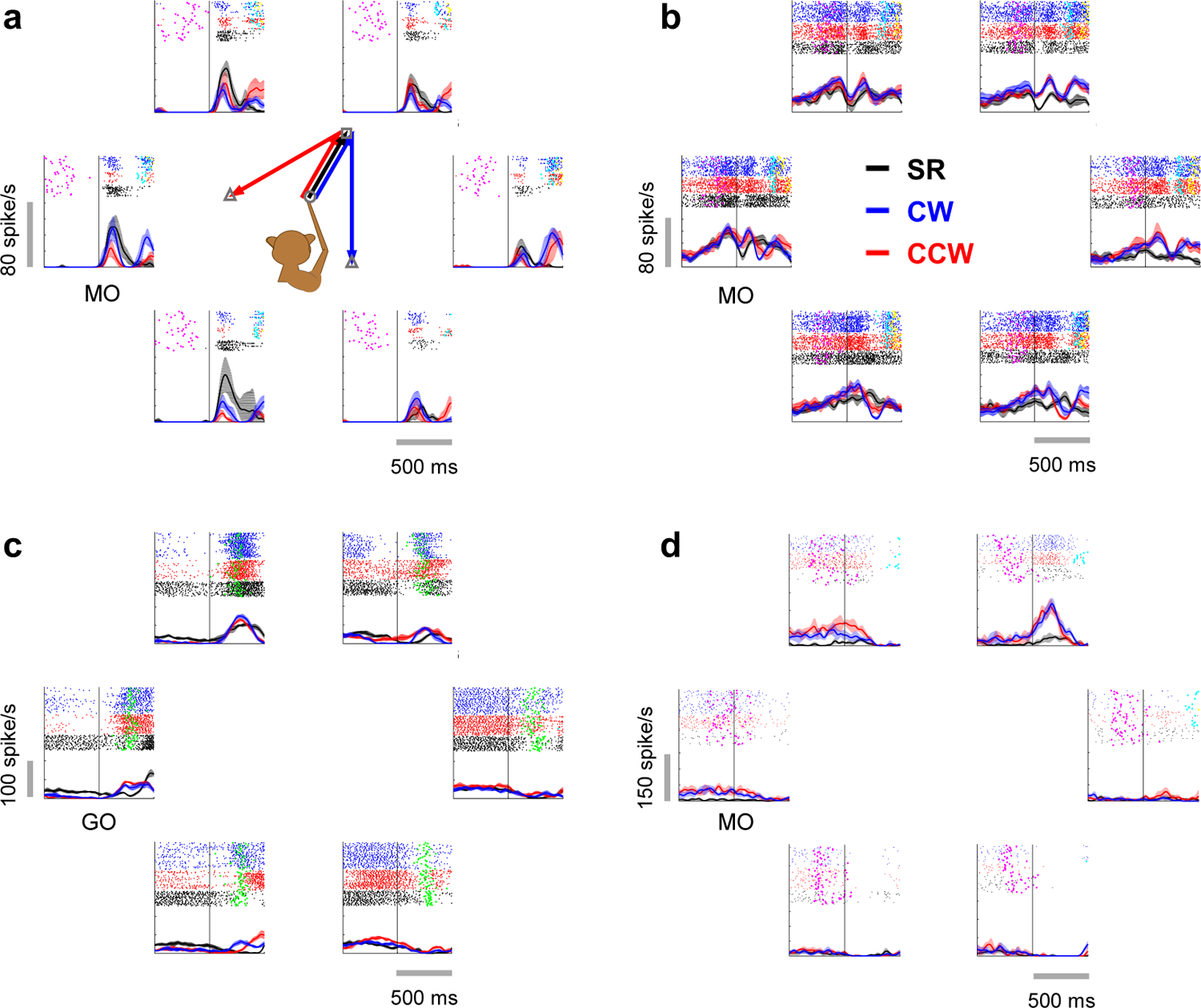
Examples of cells in motor cortex showing heterogeneous firing patterns. In each panel (a-d), the six subplots show PSTHs of the same neuron in three conditions with 1st reach toward the corresponding location (e.g., the upper-right subplot denotes the 1st reach to 60°). Rasters are plotted at the top of each PSTH (20-ms SD Gaussian kernel). Spike trains in SR (black line), CW (blue line), and CCW (red line) trials are aligned to the 1st/only movement onset (MO) in a, b, d, but aligned to GO-cue in c. Time of GO (magenta dots), MO (green dots), the 2nd movement onset (MO2, cyan dots), and the 2nd movement end (yellow dots) are presented in the rasters.

We further examined the proportion of neurons with sequence selectivity in three periods: preparatory (600 ms before GO), pre-movement (200 ms before MO), and peri-movement period (200 ms before ME). Among the 322 neurons recorded by single-electrodes, 52% exhibited significantly different firing rates for SR and DR in the preparatory period (Wilcoxon rank sum test, *p*<0.05). This proportion increased to 68% in the pre-movement period, and then to 84% in the peri-movement period (Wilcoxon rank sum test, *p*<0.05). As for the comparison between CW and CCW trials, 30%, 48%, and 72% of neurons showed significant differences during the preparatory, pre-movement, and peri-movement periods, respectively (Wilcoxon rank sum test, *p*<0.05). For the 202 array-recorded neurons, 80%, 89%, and 97% were significantly tuned to sequence during preparatory, pre-movement, and peri-movement periods, respectively (Wilcoxon rank sum test, p<0.05). In comparing CW and CCW trials, the proportions were 48%, 68%, and 87% during the preparatory, pre-movement, and peri-movement periods, respectively (Wilcoxon rank sum test, *p*<0.05). These considerable proportions reveal a substantial sequence-selectivity in the motor cortex.

### Additive *vs.* multiplicative joint coding

The above results show single-neuron responses related to reaching sequences. However, whether such sequence-related response results from joint coding or parallel coding is the next question.

Then, based on the directional tuning function:

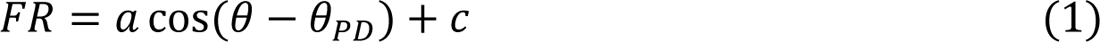

where θ is the movement direction, θ_PD_ is the PD, *a* and *c* denote regression coefficients; we developed two fitting models.

For parallel coding, the sequence-related difference comes from the overlap of two independent tuning components. For this kind of model, sequential modulation is a parallel process resulting from the preparation of the 2nd movement while the 1st movement still is in flight, as pointed out by Ames et al. ^28^. Here, we focused on directional tuning alone, and defined an ‘additive model’ as follows:

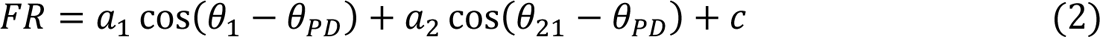

where *FR* is neuronal firing rate, θ_1_is the movement direction of the 1st reach, θ_21_ is the 2nd movement direction starting from the 1st reaching endpoint, that is, in execution coordinates, since the regression result (Fig. S2) indicates that the 2nd reach is predominately conveyed in execution coordinates (movement direction) rather than visual coordinates (target location). θ_PD_ represents the PD, *a*_1_and *a*_2_ are coefficients, and *c* is the baseline firing rate. For simplicity, we assumed the PD to be consistent for both terms at the same time.

However, since the visual targets in our task were presented simultaneously, rather than sequentially as in many previous studies ^9, 15, 16, 28^, the monkeys were more likely to prepare the entire reaching sequence beforehand ^24, 29^. In this case, the different responses in DR might not simply result from the overlap of the ‘preparation-execution’, but from interaction between the tuning components corresponding to two reaches. Therefore, this raises the possibility of joint coding, for which an interactive term is essential. For computational convenience, and as inspired by a previous study suggesting that hand speed may act as a ‘gain field’ to the directional cosine tuning function ^30^, we propose a ‘multiplicative model’ to depict the potential nonlinear gain-modulation between both elemental movements:

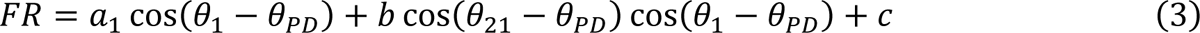

where *b* is a coefficient and other notations as in Eq. 2. If we set Δθ = (θ_21_ − θ_1_)/2, then the multiplicative term in Eq.3 can be transformed into a summation form that includes a doubled frequency (Eq. 4).

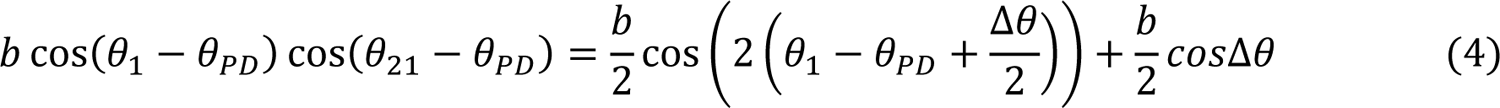

To further examine the interaction between element movements and to avoid overfitting in the regression analysis, in addition to the standard version of the paradigm described in Results (Fig. 1a), we trained monkey C to perform an extended version of the task with multi-direction, in which the angle between the square and triangle could be 60°or 120°in both CW and CCW directions as well as 180°. This multi-direction task has 36 conditions in total (six SR and 30 DR).

We tested these two possibilities on condition-averaged normalized firing rates with a 200-ms sliding window ^31^. The fitting results of an example neuron are shown in Fig. 3, in comparison with its actual PSTHs. This neuron obviously had a sequence-related mixed selectivity, because its peri-movement activity varied with different subsequent movements, and the preparatory activity was also condition-dependent, though with small variation. The response reconstructed by the additive model (Eq. 2) reproduced the peri-movement firing pattern, but it did not capture the sequence-specific modulation during preparation. In contrast, the multiplicative model (Eq. 3) better captured neural activity during the preparatory period, while losing that during the peri-movement period. In Fig. 4, we plotted directional tuning curves of the same example cell with its actual firing rates (Fig. 4, left panel), along with reconstructed firing rates by additive (Fig. 4, middle panel) or multiplicative (Fig. 4, right panel) models. The real firing rate for plotting and fitting was normalized and averaged around MO (−100∼100 ms to MO, peri-MO) and around ME (100∼300 ms to MO, peri-ME), respectively. For peri-MO (Fig. 4a), the neural tuning curves consist mostly of two peaks and were only replicated by the tuning curves of the multiplicative model. This was not accidental, because frequency doubling is a corollary of the product of two trigonometric functions (Eq. 4). For peri-ME (Fig. 4b), PD shifted with conditions in data, and only the additive model yielded a similar outcome. These results suggest that different coding rules cause distinctly different firing patterns. The multiplicative interaction contributes to the period changing, whereas the additive relation can easily lead to PD shifts while retaining the periodic identity. Comparing two epochs, the two coding possibilities could co-exist and might alternate.

**Figure 3.**
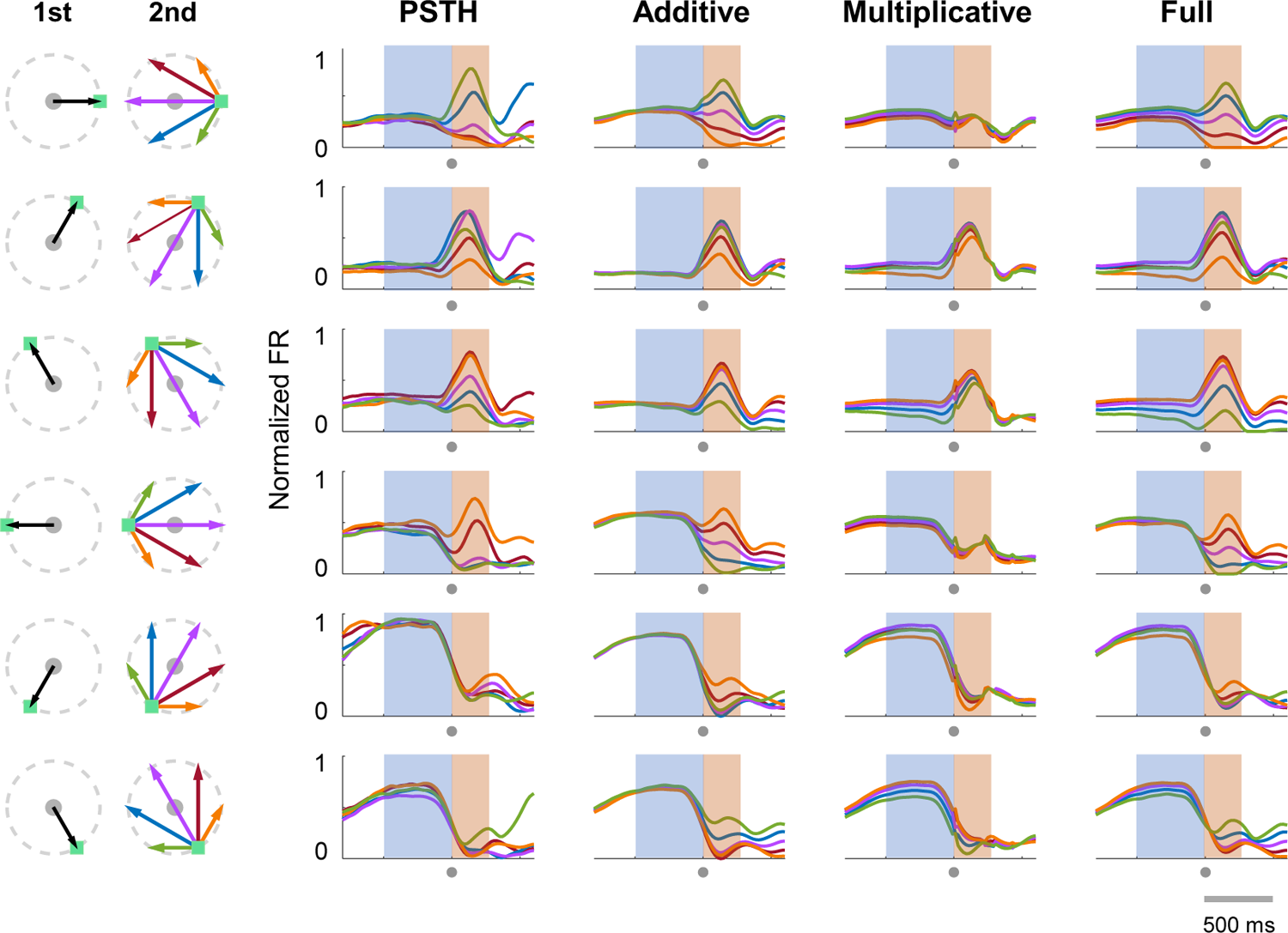
Model fitting of an example neuron. Each row shows conditions with the same 1st reach (black arrow); the 2nd reach is plotted in different colors (CW 60° in green, CW 120° in blue, 180° in purple, CCW 120° in red, CCW 60° in orange; here angle is according to the target locations in cue period). Four columns left to right are: Normalized data PSTHs; normalized firing rate reconstructed by the addictive model, the multiplicative model, and the full model, respectively. All activity is aligned to MO (marked by the gray dots under timeline, time window is −800 ∼ 600 ms to MO).

**Figure 4.**
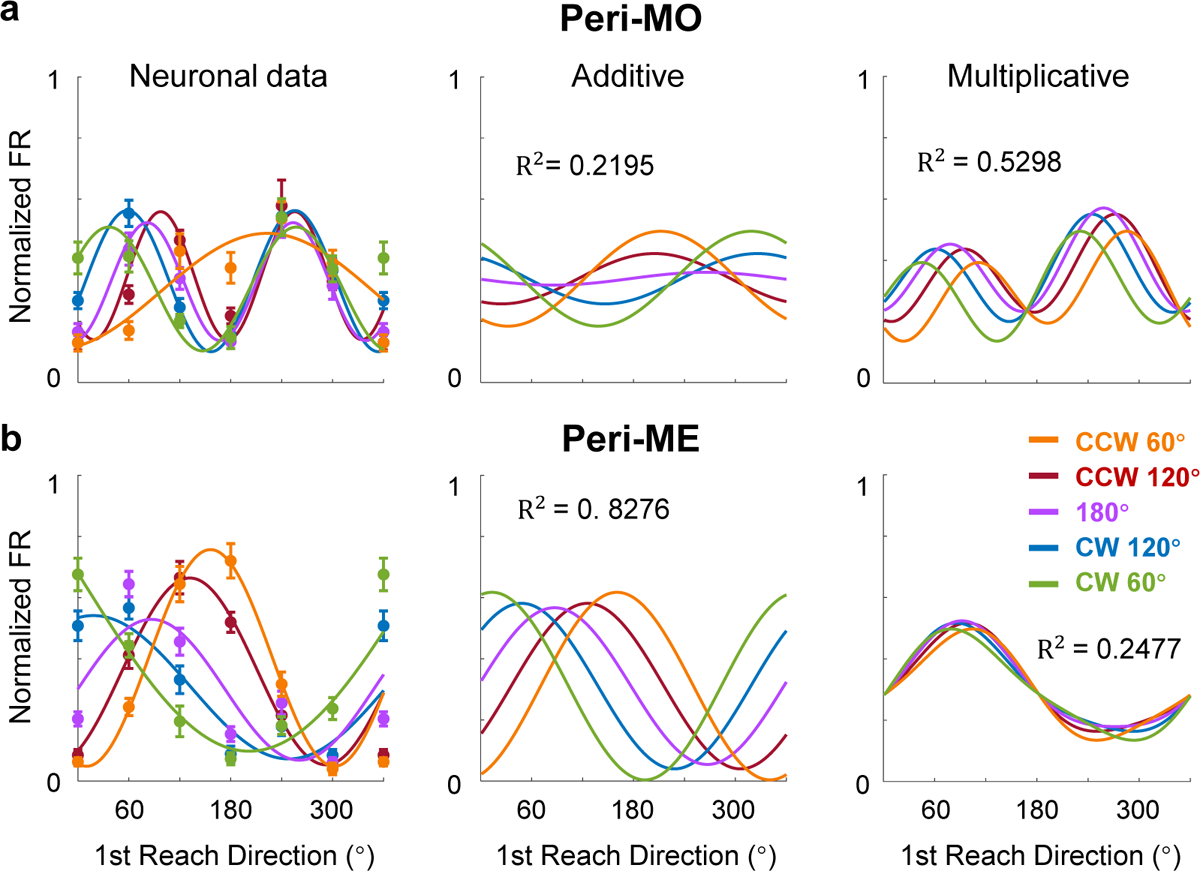
Joint tunings of the example neuron around movement onset and end. **a.** Directional tuning curves of the example cell in Fig. 3 were plotted around MO (−100∼100 ms to MO, peri-MO). Left: Normalized firing rate in DR were trial-averaged and plotted in corresponding condition colors. Tuning curves were fitted by Fourier expansion separately. Middle: Tuning curves of firing rates reconstructed by the addictive model. Right: Tuning curves of firing rates reconstructed by the multiplicative model. R^2^ showed the goodness-of-fit of the model tuning curve. **b.** Similar with **a**, directional tuning curves around ME (100∼300ms to MO, peri-ME).

To further investigate the temporal dynamics of joint-coding rules, we proposed a ‘full model’ to combine the two modulation forms:

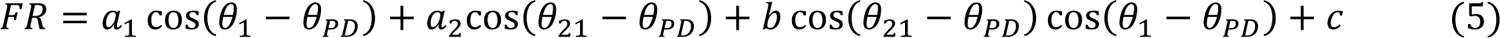

where descriptions of notations are the same as in Eq. 2 and Eq. 3, defining *a*_1_as the 1st reach weight, *a*_2_as the additive weight, and *b* as the multiplicative weight. The fluctuation of the regression coefficients (*a*_1_, *a*_2_, and *b*) reflects the time-varying contribution of the corresponding terms, thus enabling the full model to profile the transition of coded objects.

We compared the goodness-of-fit of the full model with that of the additive model, the multiplicative model, and a single cosine model (Eq.1, 1st reach direction), by the standard of the population-averaged adjusted R^2^ (For the statistical method to compensate for the difference in numbers of parameters between the full model and other models, see Methods) for M1 and PMd, respectively (Fig. 5, array data from monkey C). The full model performed best in both areas; it was also able to describe the tuning property of the example neuron throughout the whole trial (Fig. 3, Full). Remarkably, the overall trend and preference for the multiplicative or additive model varied by brain areas. For M1 neurons (n=118), the goodness-of-fit for all models gradually increased during preparation, and the multiplicative model was significantly better than the additive model at MO (two-tailed Wilcoxon signed rank test, p=1.2e-05). Nevertheless, the additive model performed better after MO. Similar results were found in M1 data in all monkeys (Fig. S3, single-electrode recording from monkeys B and C, and array data from monkey G). The effect size *r* (see Methods) also indicates there is a small to medium effect for multiplicative model during preparatory period for each monkey (Fig. S4). For PMd neurons (n=40), the adjusted R^2^ remained stable during preparation. No significant difference was found between two models before MO (two-tailed Wilcoxon signed rank test, p=0.06). It seems that the transition from multiplicative to additive coding was different in M1 and PMd.

**Figure 5.**
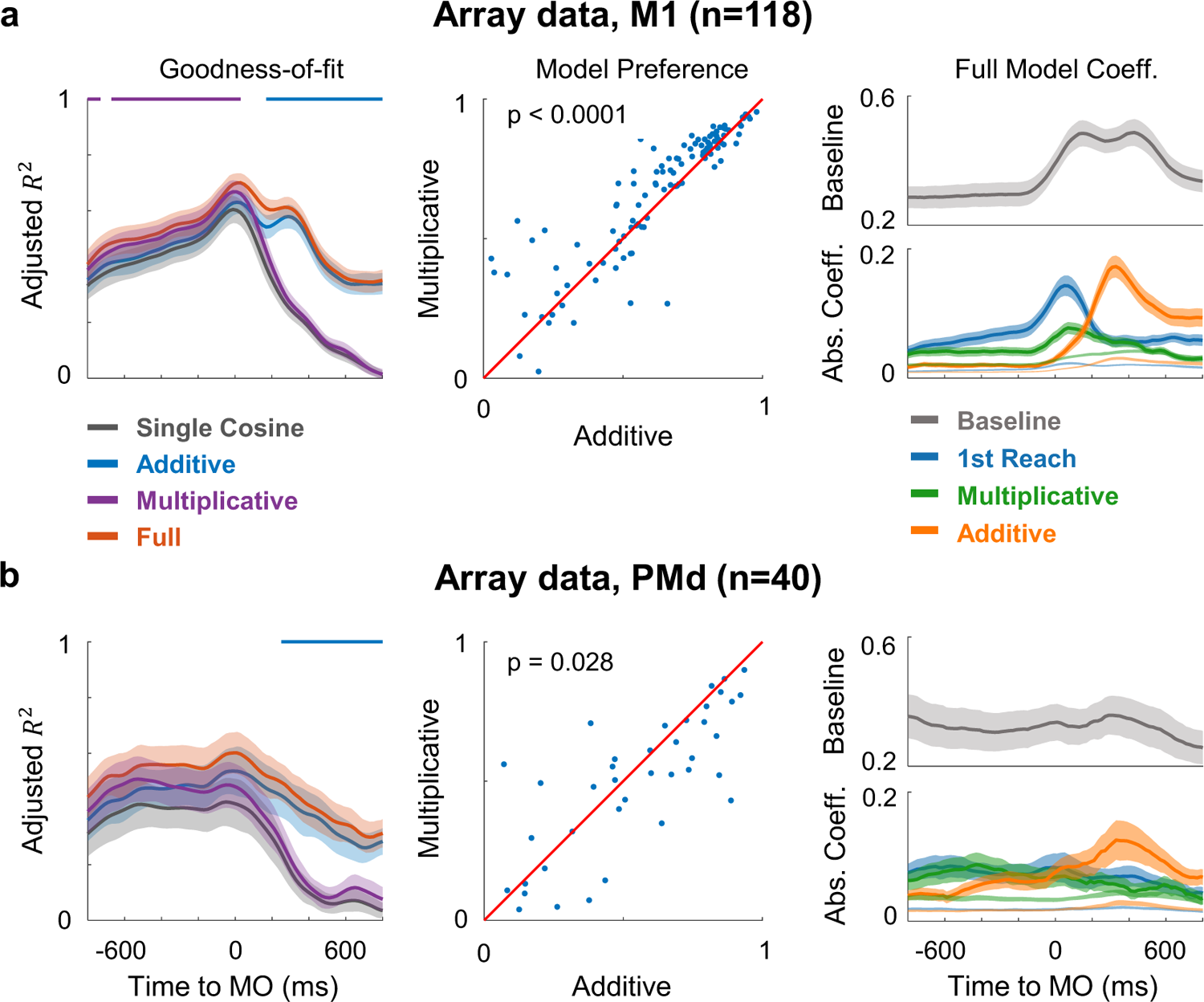
Dynamics of goodness-of-fit and coefficient. **a.** Results of regression on M1 neurons in array dataset from monkey C are illustrated at the population level. Left: Goodness-of-fit was evaluated with averaged adjusted R^2^ for all fitting models in a 200-ms sliding window (with twice standard error in shade). The upper line showed the significance (p<0.0005) of comparison between performance of multiplicative (purple line) and additive (blue line) model. Middle: Scatters compared the goodness-of-fit at MO (−100∼100 ms to MO) between the multiplicative and additive models, each dot represents the result of a neuron. Right: Absolute value of each coefficient is averaged across neurons (with twice standard error in shade), the temporal dynamics of which shows the contribution of terms. The coefficient weight of permutation test was plotted in light shade as the chance level. **b.** Similar with A, results of regression on PMd neurons in array.

To scrutinize the changing encoding pattern, we plotted the averaged absolute coefficients of the full model across time (Fig. 5, right panel). For M1 neurons, the weights of the 1st reach and the multiplicative term ramped up over the chance level (given by a permutation test, see Methods) during preparation, whereas the additive weight remained at the chance level in preparation and mainly increased after MO. This contemporaneous activation of coefficients was similar to the situation in prefrontal cortex where neurons were modulated by both direction and sequence ^32–34^. Similar dynamics were found in all monkeys (Fig. S3), suggesting a common transition from a gain-modulation interplay during motor preparation to a concurrent coding during motor execution. This concurrence has been reported by the previous study ^27^. For PMd neurons, the overall encoding process was essentially consistent with that in M1: the 1st movement → the multiplicative term → the additive term. However, in PMd, these three components made comparable contributions in the preparation period, and there was no obvious peak of the 1st reach and multiplicative coefficients. The onset times for the increase of the additive coefficient, and the decrease of the 1st reach and multiplicative coefficients, were much earlier in PMd than in M1, implying that PMd takes precedent in coding of the 2nd reach.

So far, we have analyzed the linear and nonlinear components comprised in neural encoding for double-reach and their interchangeable predominance. The multiplicative joint coding, revealed by the multiplicative model and validated by the multiplicative weight in the full model, now becomes a key concern because it would be apparently a unique signature of continuous motor sequences.

### Multiplicative coding embodied in initial states

According to our regression analyses, the multiplication of the tunings corresponding to the 1st and 2nd reaches could be intrinsic in sequence-related preparatory activity. From the dynamical systems perspective, preparatory activity would be set to a subspace optimal as initial states to trigger motor generation ^35^. We expected a spatially inclusive distribution of initial states to accord with mathematical multiplication.

To verify this hypothesis, we performed a supervised dimensionality reduction procedure. Firstly, principal component analysis (PCA) was applied to the preparatory neural activity during a period of 600 ms before GO. Next, the Fisher’s linear discriminant analysis (LDA) was utilized to find the optimal discriminant projection in accordance with tagged conditions ^36^. In this PCA-LDA analysis, selected principal components from PCA (the number was chosen by cross-validation) were applied to LDA. We first analyzed neural activity in SR trials and built an SR subspace. Neural states clustered by conditions, as visualized in the 2-d projections found by LDA (Fig. 6a). Then, we projected both DR and SR data onto the resulted space and found that neural states of both DR and SR trials clustered according to their 1st or only reach direction. This suggests that despite the proposed sequence modulation in preparatory activity for single neurons, the neural population preserved a linear representation for the preceding movement. However, the variance explained were higher for SR than DR (For monkey C array, variance explained of SR is 10.9%, DR is 7.8%. For monkey B, variance explained of SR is 9.0%, DR is 6.6%. For monkey C single electrode, variance explained of SR is 6.2%, DR is 5.6%. For monkey G, variance explained of SR is 31.6%, DR is 27.5%.). To neutralize the tuning for the immediate movement, we used DR trials with the same 1st reach direction alone for the PCA-LDA analysis. Therefore, relatively low-dimensional neural states grouped by the 1st reach directions, could be projected again onto dimensions maximizing the difference brought by the 2nd reach directions. The result of trials where the 1st reach direction was towards the lower-right was visualized, with trials classified into six clusters corresponding to their subsequent reach directions (Fig. 6c; subsequent reach directions are indicated by markers; ten-fold cross-validation accuracy was higher than 0.6, above the chance level for the classification of six conditions, 1/6, excluding LDA overfitting). There were great differences between SR (circles) and DR (other markers) clusters, indicating that the initial states for sequential movements were distinctive. Interestingly, in some conditions, DR trials obviously clustered in order from CW 60°to CCW 60°, and the CW and CCW states were located on both sides of the 180°states. This structural spatial distribution of LDA states supported by Mahalanobis distances (Fig. S5) may signify a condensation of subsequent movement information in the strong representation of occurrent movement. In addition, the results for other monkeys for the DR task showed a similar tendency (Fig. S6-S8).

**Figure 6.**
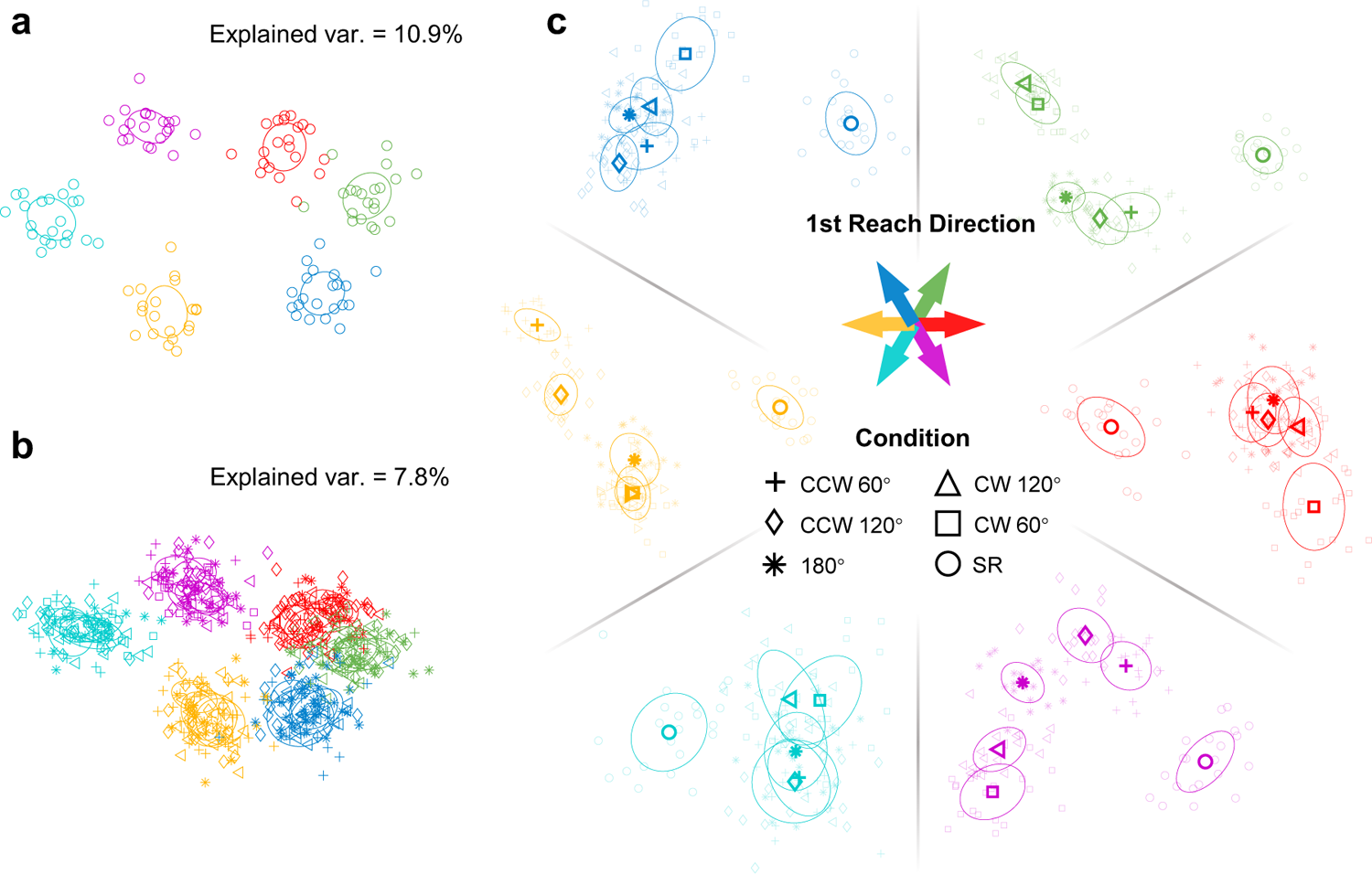
Projection of preparatory activity onto PCA-LDA resulting initial state space. **a.** Projection on SR space. Neural states of SR trials were clearly clustered according to their reaching directions. **b.** Neural states of DR trials also clustered into six groups according to their 1st reach direction when projected onto the SR space. Variance explained by the two dimensions were calculated. **c.** LDA classified neural states of trials with the same 1st reach direction into clusters grouped by 2nd reach directions, forming an initial state space for the subsequent movement. Colors indicate the 1st movement directions; DR trials are presented in the same color family of related SR trials. Markers indicate 2nd reaching direction. The ellipses show the covariance projection of related conditions.

### Multiplicative coding preserves linear readout of immediate reach

As several investigators have pointed out ^12, 21, 37, 38^, as well as PCA results suggested, the neural population preserves a reliable readout of ongoing movement direction, despite the sequence-related differences at the single-neuron level. Since nonlinear mixed selectivity is believed to form high-dimensional neural representations that guarantee the linear readout of particular parameters ^39^, we speculate that each linear readout in sequential movements benefits from multiplicative joint coding.

We checked the linear readout of immediate movement direction as population vectors (PV) in data. The PV pointed to the immediate reach direction before MO in DR trials as expected (Fig. S9). To figure out the impact of the multiplicative or additive joint coding on PV, we adopted a proved simulation method ^40^ to obtain surrogate data corresponding to the cosine, additive, and multiplicative models (see Methods). Each dataset consisted of 200 model neurons with activity in an epoch of 600 ms from preparatory activity until the 1st reach end. Those additive and multiplicative neurons were regulated by a fixed 2nd reach direction as well. We present the responses of three example model neurons with the same θ_PD_ in Fig. 7a. Obviously, the direction inducing the highest firing rate changed in additive and multiplicative neurons, compared to the ‘single cosine’ neurons, resulting in a modulated tunning curve (Fig.7b). We used the original θ_PD_for the calculation of PV. Interestingly, PVs in the multiplicative DR dataset correctly and stably pointed to the immediate reach direction as in the SR condition, whereas PVs in the additive DR dataset deviated from the desired direction (Fig.7c). These simulations show that multiplicative joint coding can preserve a robust linear readout of immediate reach direction, even containing subsequent reach directions.

**Figure 7.**
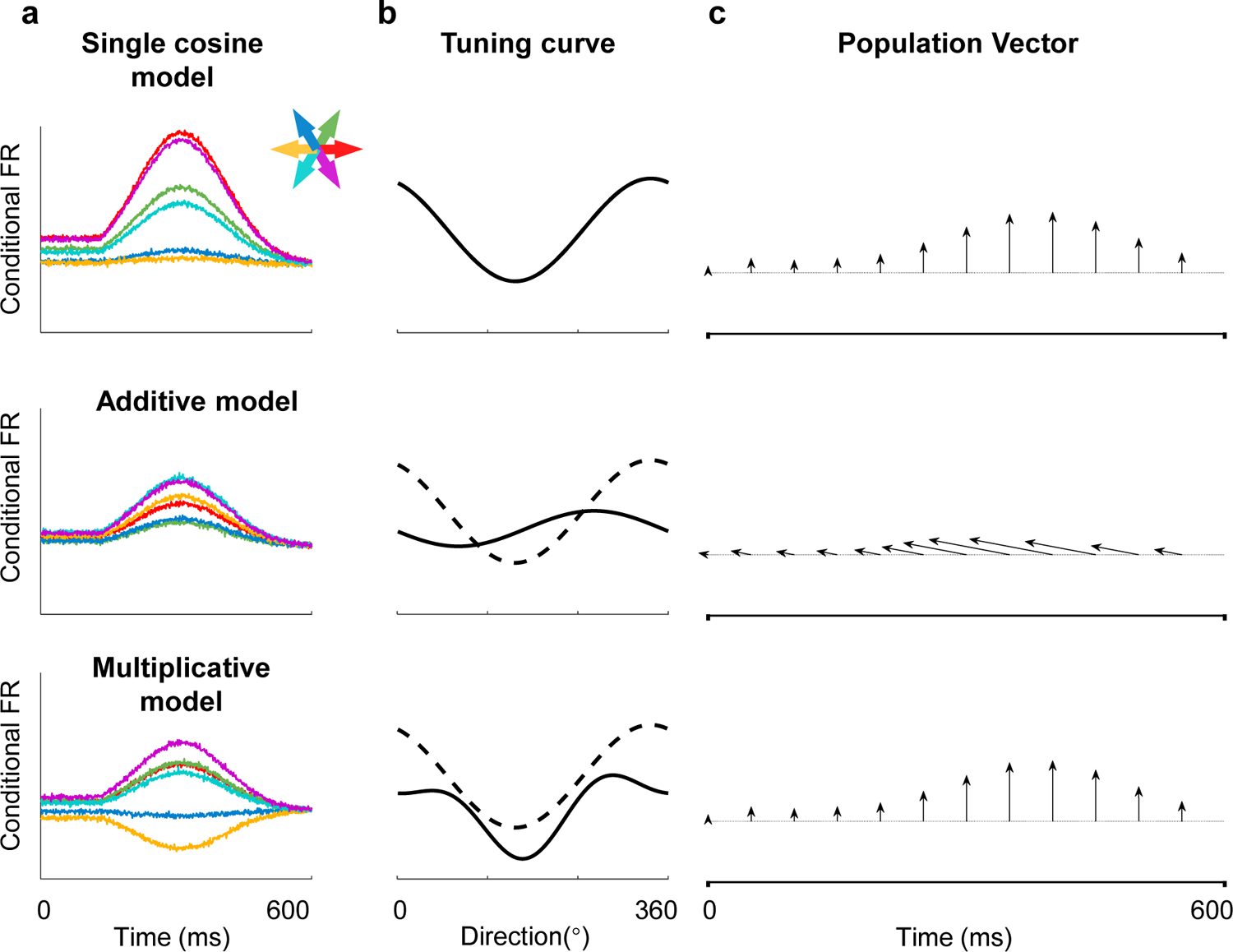
Simulation of neural tunings on population vector during single and double reach. **a.** Example neurons of three simulated datasets. Averaged firing rates of different conditions (1st reach directions) are shown in corresponding colors. These three example model neurons were simulated according to the single cosine, the additive, and the multiplicative models with the same preferred direction θ_PD_. **b.** Directional tunning curves with (solid line) and without (dash line) modulation. **c.** Population vectors of three simulated datasets. Population vectors were calculated every 50 ms. The correct reaching direction is upward. The population vector of multiplicative dataset pointed in the same direction as PV of single cosine dataset, while the PVs of additive dataset shift away from the desired reaching direction.

### Multiplicative joint coding emerged in recurrent neural network (RNN) generating motor sequence

Due to their flexibility and time-varying characteristics, RNNs are increasingly welcomed as models matching a dynamical system ^41–43^. To find out whether a dynamical system can also capture the subtle joint-coding rule found in the motor cortex, we trained an RNN model to perform the double-reach task.

The three-layer RNN received the movement direction of two reaches as input (Fig. 8a). The input signals were presented simultaneously, though instructing sequential actions. In contrast to previous work in which RNNs were instructed to generate velocity ^40^ or EMG ^44^, our model was required to produce PV. This design was preferred for these reasons: first, the variables related to actual movement, like velocity and EMG, have to lag behind the neural activity due to transmission delay from cortex to muscle. In contrast, PV could be real-time, and thus reflect more temporal features; also, this design is consistent with our hypothesis that multiplicative joint coding benefit linear readout of movements (Fig. 7).

**Figure 8.**
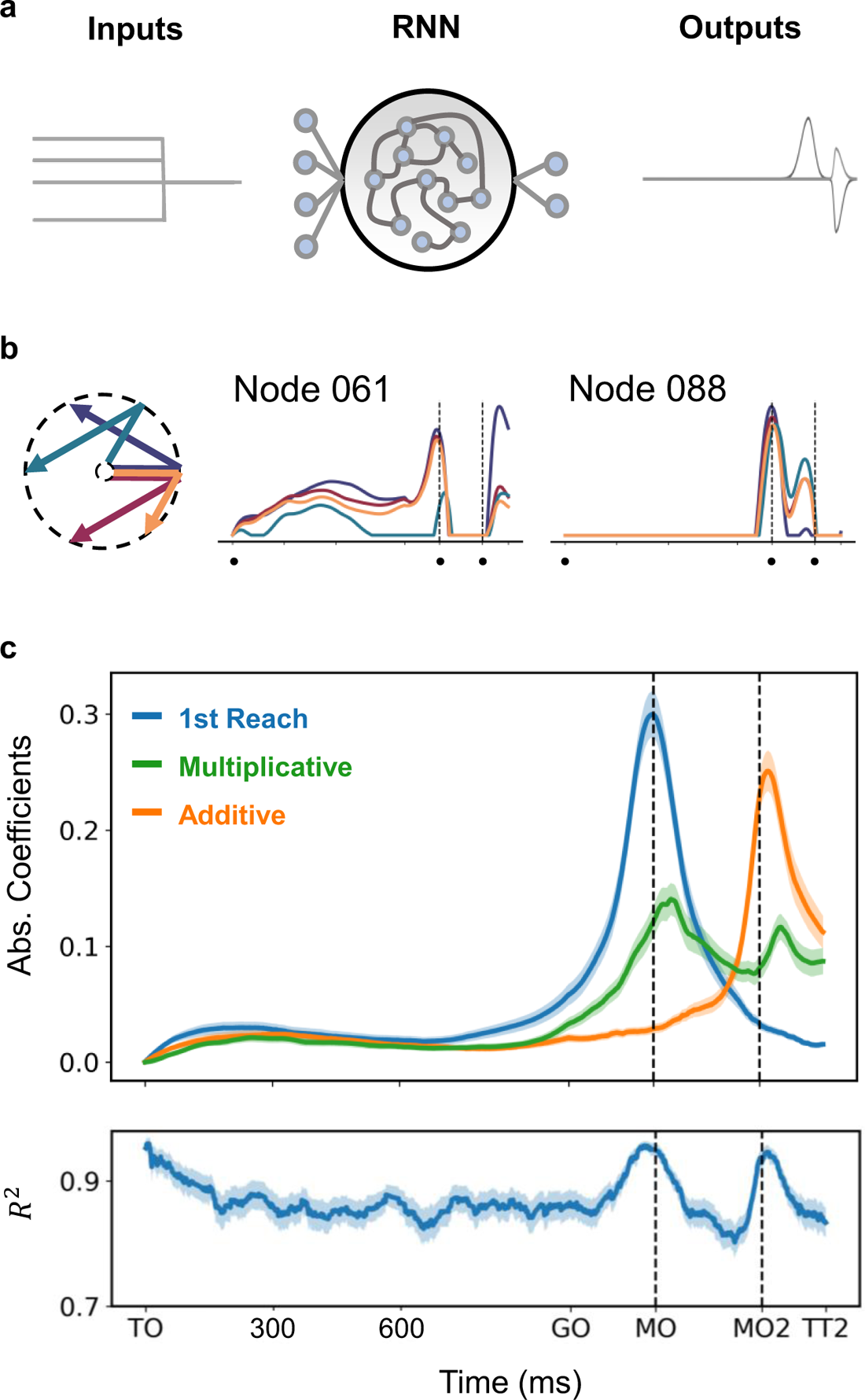
Results of an RNN model. **a.** Schematic of the RNN model. The RNN model consisted of an input layer, a hidden layer, and an output layer. The input layer received signal for position of two targets simultaneously, while the output layer produced population vector (PV), whose magnitude reflects the degree of movement tendency for neural population in the corresponding direction. **b.** Response of two example nodes under four conditions. The selected conditions are represented in different colors as shown in left. The black dots denote target on (TO), the 1st movement onset (MO), and the 2nd movement onset (MO2), respectively. Most model nodes show temporally similar responses to real neurons. **c.** Full model fitting result for RNN nodes. It turned out that the temporal dynamics of the terms in this RNN model are comparable to those in real neurons. The R^2^ was calculated across nodes. The error bar in this panel was plotted according to twice standard error. Time markers are ticked as: target on (TO), go-cue on (GO), the 1st movement onset (MO), and the 2nd movement onset (MO2).

The trained RNN performed well (for training set R^2^=0.9711±0.0044, for validation R^2^=0.8976±0.1070, mean±SD; see Methods). Model nodes exhibited comparable temporal dynamics with real neurons recorded in the present study. Here we exhibit two example nodes under four specific conditions (Fig. 8b). For the node 088, the two bumps of its response indicate that it is closely related to the ongoing movement, which is typical for neurons in M1. The response of the node 061 seems more complex, as the augment around MO does not occur in all conditions. Interestingly, this node appears to have ‘direction selectivity’, the only exceptive movement direction for the 1st reach (in cyan) induces obviously distinguished response. Moreover, its responses under different conditions retain distinct during preparatory period.

Observing richer preparatory dynamics than expected, we wondered whether the temporal dynamics of components corresponding to different movement courses were consistent between model and neural data. Therefore, we tested the ‘full model’ fitting on nodes of our model. As shown in Fig. 8c, the profile of regression coefficients of model nodes largely resembles that of real data (Fig. 5a, right, Fréchet distance = 0.47, see Methods). The weight of the 1st reach peaks at MO and decays afterwards. The weight of the additive term, which relates to the 2nd reach, reaches its apex around MO2 with a slightly smaller magnitude. During the preparation, the weight of the multiplicative term fluctuates, but maintains a considerable influence. This suggests that the proposed multiplicative joint coding for sequential movement, here a double reach, also emerges in a dynamical system.

## Discussion

In order to understand how the motor cortex generates motor programs for consecutive arm movement sequences, we recorded neuronal activity when monkeys performed double-reach directed at simultaneously cued memorized targets. We found that pre-movement activity carries sequence information in a heterogeneous manner. Regression analysis shows that neuronal tuning to 1st and 2nd reaches can be well explained by multiplicative and additive models in the preparatory and execution periods, respectively. Dimensionality reduction analysis demonstrates that neural states during preparation sub-clustered according to the 2nd reach within the optimal subspaces of the 1st reach. Simulation via model neurons points out the merit of multiplicative joint coding in maintaining robust linear readout for the ongoing movement direction. An RNN model trained for double-reach task can simulate the real encoding properties, which are marked by conspicuous nonlinearity. Taken together, these results suggest that primate motor cortex is profoundly involved in forming plans for multi-step movements. In addition, the transition between the newfound multiplicative joint coding and overlapped independent coding hints at a shifting neural encoding mechanism for motor sequences.

Previous studies have revealed that motor cortex not only carries information regarding upcoming movements, but also reflects sensory and cognitive factors during both preparation and execution periods ^6, 9, 12, 16, 17^. Nevertheless, how this ‘sequence selective’ response reflects motor sequence has not yet been answered. A recent work following the dynamical systems perspective found that ‘the preparatory subspace was occupied twice, once before each reach’ thus suggested that each of movement elements were encoded independently in the motor cortex rather than holistically. However, if individual movements were independently planned, the reaching error should accumulate, which has not yet been observed ^45^. Furthermore, it was demonstrated that the holistic planning might enhance motor learning, but such effect would not occur when different follow-throughs were rehearsed individually ^46^. This finding strongly suggests that sequential planning is associated with special neural states in preparation, in accordance with our findings. Also, unlike in the parietal cortex, neuronal activity in the motor cortex exhibits strong heterogeneity ^47^, which often comes from mixed selectivity of behavioral parameters and tuning dynamics ^30, 48–50^. Given these considerations and our results, we propose that elements in a consecutive movement sequence should be interactively planned in a spatio-temporally coordinated manner beforehand.

As one of the cortical regions carrying much information regarding movement timing ^10^ and kinematics ^11^, motor cortex presumably participates in encompassing and coordinating sequence components. In the present study, both reaching targets were turned off 400-800 ms before GO, encouraging the monkeys to plan the whole reaching sequence in the preparatory period. Our results revealed that neurons tended to jointly encode double-reach in a nonlinear multiplicative manner during preparation. The multiplicative model’s performance degraded after MO, perhaps because joint coding mainly exists during preparation, but lies in the null-space during execution. Also, as a reaching sequence is decomposed into motor elements, the lack of an additive term makes it incapable of capturing the parallel components after MO ^27^. The concept of the multiplicative model originated from gain modulation ^51, 52^, and a work regarding the question of whether the neural response was constructed with nonlinear interactions between parameters, rather than their linear combination ^30^. In the case of sequential movements, this issue becomes whether sequential elements are planned conjunctively or independently. As the primary nonlinear interaction, multiplication is a common form of gain modulation that has been widely found in mixed selectivity ^51, 52^. This coding manner can provide new dimensions for motor preparation and learning ^53^, and according to our simulation, it can also consolidate the linear readout for impending movements. Because such mixed selectivity of parameters enlarges the neural space encoded by a certain number of neurons ^36, 39, 53^, the dimensionalities induced by multiplicative coding may perform as the null space of impending movement.

Although our analyses of joint coding are based on directional tuning, we did not mean to imply that the motor cortex exclusively encodes movement direction. Rather, we treated the directional tuning as a marker of interaction, rooted in the heterogeneous neuronal response. Since the motor cortex is recognized to play a straightforward role in generating descending command for muscle activity production ^54, 55^, future studies should also take into account muscle activity to explain how joint coding benefits the generation of compound double reaches from a dynamical systems perspective ^44^. However, it is a limitation of the present study that sEMG data were not sufficient to explore this issue.

Regarding joint coding embodied in motor cortex as a key signature to encompass movement elements in the planning of consecutive sequences, we are not claiming that it seeds a neural dynamical system that can autonomously generate the entire motor sequence. Instead, sequential behavior emerges from a large brain network, including parietal-frontal circuits ^9, 56^ and subcortical areas like the thalamus and basal ganglia ^57^. Remarkably, our results of PMd, which are so different from those of M1, have already indicated diverse functions and coding characteristics of different cortical regions. Now that the dynamical evolution in the motor cortex necessarily relies on external inputs from other brain areas ^58^, an intriguing question is how intrinsic dynamics and external inputs interplay to generate a motor sequence, including the role of the proposed joint coding in the motor cortex. To go further, collective studies across multiple brain regions and experimental interventions are needed.

## Acknowledgments

We thank Y. Guo, J. Malpeli, Q. Wang, C. Zheng, and R. Zheng for helpful comments and discussions; C. Guan and L. Wang for veterinary assistance; P. Ding for administrative support.

## Author Contributions

T. Wang, Y. Zhang, and H. Cui designed the experiment, T. Wang and Y. Zhang collected the data, T. Wang analyzed the data, Y. Chen built computational model, T. Wang, Y. Chen, and H. Cui prepared the manuscript.

## Methods

### Experimental preparation

Three male rhesus macaques (monkey B, C, and G, *Macaca mulatta*, 5-9 kg) were trained to perform a cohesive double reach (Fig. 1a). In each session, the monkey sat in a custom-designed primate chair. Stimuli were backprojected onto a vertical touch screen (Elo Touchsystems, 19”; sampling at 100 Hz, spatial resolution <0.1 mm) ∼30 cm in front of the monkey. In the recording sessions using microelectrode arrays (Utah array, Blackrock), hand position was monitored optically via reflective markers attached to the wrist (Vicon Inc.), besides, acceleration and surface electromyography (sEMG) were recorded via a wireless sensor (Delsys Trigno Lab) attached to the targeted muscles. All procedures were in accordance with NIH guidelines and were approved by the Institutional Animal Care and Use Committee (IACUC) of Institute of Neuroscience, CAS.

### Behavioral task

In addition to the standard version of the paradigm described in Results (Fig. 1a), to further examine the interaction between movement elements, we trained monkey C to perform an extended version of the task with multi-direction, in which the angle between the square and triangle could be 60°or 120°in both CW and CCW directions as well as 180°. This multi-direction task has 36 conditions in total (six SR and 30 DR).

### Data collection and analysis

For single-electrode recording, monkeys B and C were implanted with a standard recording cylinder (diameter = 19 mm) located over M1 and caudal PMd in the left hemisphere, guided by pre-scanned MRI and stereotactic coordinates. Recording sites are shown in Fig. S1. Recordings were made using glass-coated tungsten electrodes (AlphaOmega, ∼1.5 MΩ impedance at 1 kHz). Activity was recorded online by an AlphaOmega Lab SNR system, sampled at 44 kHz. After recordings, raw data were sorted offline according to an online template by Spike2 (Spike2 7.15, CED). For multi-electrode recording, monkey G and C, respectively, were implanted with a 96-channel and two 128-channel Utah microelectrode arrays (Blackrock Microsystems, Salt Lake City, UT) in the motor cortex of the right hemisphere (Fig. S1). Recording sites were located using MRI and cortex surface features. Array recorded raw data were sorted offline by Wave_clus ^59^.All monkeys were restricted to using the hand contralateral to the recorded hemisphere when performing the task. Data from monkey C were first obtained with a single microelectrode, and subsequently from an array in the other hemisphere with a switch of hands.

In total, we collected 279 and 117 well-isolated units from monkey B and C through single-electrode recording, respectively. Among these, 224 units from monkey B and 98 from monkey C with significant directional preference (One-way ANOVA, p<0.05) in single reach were chosen for further analysis. For multi-electrode recording, we collected 252 and 63 well-isolated units from monkey C and G, respectively. Among these, 158 units from monkey C and 44 from monkey G with significant directional preference (One-way ANOVA, p<0.05) were used. The selected neurons formed a 3-dimensional *NKT* (*N*: neuron number, *K*: trial number, and *T*: spike time) dataset for regression and state-space analysis.

### Peri-stimulus time histograms (PSTHs)

For each unit, we calculated its PSTHs with time aligned to event markers such as the GO signal, the 1st/only movement onset (MO), the 1st/only movement end (ME), and the 2nd movement onset (MO2). We defined MO as the moment when the monkey’s hand left the touch screen and ME as the time when monkey’s hand touched the target on the screen. All firing rates were smoothed with a Gaussian kernel (SD = 20 ms). The mean standard error (mean SE) of firing rate was estimated from 10 bootstrap samples.

### Regression

We adopted the directional tuning model ^1, 30^ to fit neural responses in the double-reach task. We fitted the normalized condition-averaged firing rates in a 200-ms sliding window with 20-ms step (using Matlab function ‘fit’ and ‘fitnlm’). First, we fitted the double-reach data as follow:

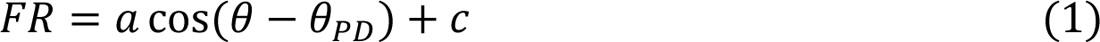

where θ is the movement direction, θ_PD_is the PD, *a* and *c* denote regression coefficients. Both the 1st and the 2nd reach direction were used for regression to see which direction is better represented at that time bin (Fig. S2). Then we regressed double-reach data with the following models:

Additive model:

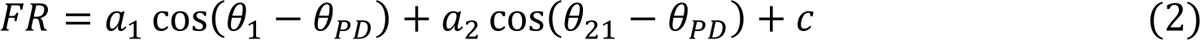

Multiplicative model:

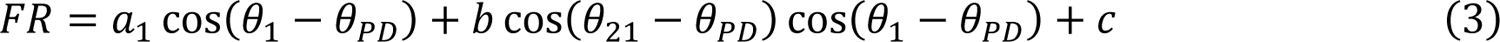

Full model:

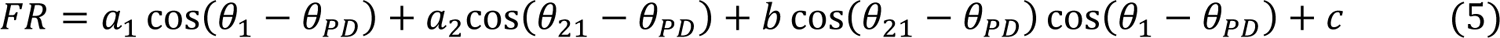

where *a*_1_, *a*_2_, *b*, *c* are regression coefficients, θ_1_ is the 1st movement direction, θ_21_ is 2nd movement direction from the 1st reach endpoint, θ_PD_ is preferred direction.

Note that both the additive and multiplicative model have four coefficients while full model has five. To compensate for this difference, we use the adjusted R^2^ rather than actual R^2^.

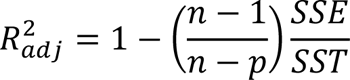

where SSE is the sum of squared error, SST is the sum of squared total, *n* is the number of observations, and *p* is the number of regression coefficients. Because actual R^2^ likely increases with added predictor variables in the regression model, the adjusted R^2^ adjusts for the number of predictor variables in the model. This makes it more useful for comparing models with a different number of predictors.

We also compared the goodness-of-fit between the multiplicative and additive models using the Wilcoxon signed rank test. We plot a line (purple for the multiplicative model, blue for the additive model) when one is significantly (p<0.0005) better than the other.

In addition, we calculated the effect size *γ* = *Z*/√*n* using function “wilcoxonPairedR” in package “rcompanion” of R (Mangiafico, S.S. 2016. Summary and Analysis of Extension Program Evaluation in R, version 1.19.10. rcompanion.org/handbook/). The *γ* value could be interpretated as small effect in 0.1-0.4, medium effect in 0.4-0.6, and large effect ≥ 0.6.

To get the chance levels of each coefficient and to reflect the effect of modulation, we performed a permutation test with 1000 repetitions separately for the coefficient of the 1st reach, multiplicative term, and additive term in reference of Sober and Sabes ^60^.

### PCA-LDA analysis for neural states

*NKT* datasets were used in this analysis. Neuronal firing rates were calculated with a 300-ms bin width (*T* = 2) and normalized by Z-score (MATLAB function ‘*zscore*’) to avoid bias from high firing rate neurons. *NKT* data were reshaped into *K* × *NT*, where *K* is trial number, *N* is neuron number, and *T* is bin number. After pre-processing, we ran PCA to reduce these dimensions to *K* × P. The number of PCs, P, was chosen by 10-fold cross-validation to avoid overfitting. This step also helped avoid singular matrices for LDA and reduced data noise ^36^. Then we ran LDA to project the P-dimensional matrix onto a *C*-dimensional space, where *C* is the number of trial conditions. LDA can find axes that best separate the categories. After this, we applied QR decomposition to get the orthonormal basis for the neural state space ^61^. Each trial was finally described by *C* − 1 components derived from selected neural activity. We chose the first two components covering the largest variance to plot the 2-D projection of trial data and the ellipse of covariance; each data point represented the neural state in a trial.

### Simulation of population vector in sequential reach

We adopted the simulation method of ^40^ to generate surrogate data based on single cosine, additive, and multiplicative models. Preparatory and peri-movement activity were simulated with 200 neurons in six directions. The averaged neuronal firing rate *f*_*n,c*_ for neuron *n*, in condition *c*, at time *t* is given by,

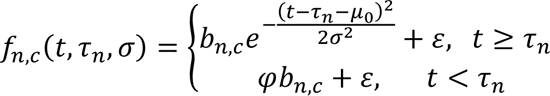

where σ is the duration parameter, *τ*_*n*_ is the response latency of each neuron (normally distributed), Φ is the preparatory activity amplitude constant fixed at 0.2, μ_0_ is constant given by μ_0_ = σ√−2lnΦ, and ε is random noise (SD=0.01). *b*_*n,c*_ is the gain for neuronal condition preference. For data of the cosine model, which is expected to mimic neuronal activity in SR trials, *b*_*n,c*_ is simply tuned to reach directions as

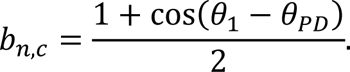

The additive surrogate data were based on the parallel coding hypothesis that sequential movements are planned independently with the overlap in the peri-movement period; *b*_*n,c*_ is given by

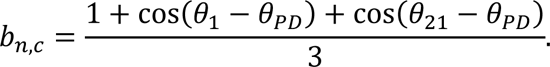

The multiplicative surrogate data were based on the gain-modulation hypothesis, the interaction of both movement directions in sequential reach contributed to the neuronal response,

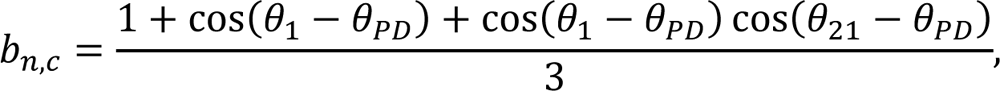

For the above definitions, θ_1_is the 1st movement direction, θ_21_is the 2nd movement direction relative to the 1st movement endpoint, and θ_PD_ is the preferred direction.

### Model training

Our RNN model was designed to simulate the situation where double reach was accomplished by a pure dynamical system. The input was movement direction for two sequential reaches, in form of 2-D coordinates [cos(θ_1_), sin(θ_1_); cos(θ_2_), sin(θ_2_)], where θ_1_ and θ_2_ represent the 1st and relative 2nd movement directions, respectively. Because the model was built to generate population vectors (PVs), we constructed ‘desired PVs’ instead of using real data for generality. The output was read out as [*γ* cos(θ), *γ* sin(θ)], where θ is the present movement direction, and *γ* reflects the intensity of integrated response for population. We used Gaussian functions to emulate the time-varying magnitude. To ensure the trend at critical time markers was similar to the actual situation, we separated the two-peak PV profile into four sections: from GO to MO, from MO to the 1st touch, from the 1st touch to MO2, and from MO2 to the 2nd touch, and spliced them together after respective optimization and normalization. We used 18 standard conditions in training, including SR conditions and DR conditions as mentioned previously. For validation, we tested 30 conditions, in which the angles between the 1st and 2nd targets were 60° and 120°in both CW and CCW directions, as well as 180°.

The nodes in the RNN model were evolved according to a standard continuous dynamical equation ^40^:

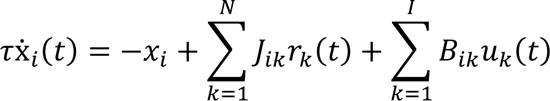

where *τ* is a time constant, *N* is the number of network nodes, and *I* is the number of the inputs. The activity of nodes is represented by *x*, whose firing rates are determined by

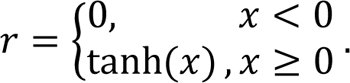

The output was read out linearly as:

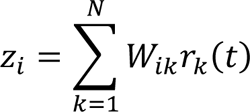

where *Z* represents the two PV readouts (*i* = 1,2). In this model, the connection weight among nodes is denoted by matrix *J*, the connectivity between hidden nodes and input *u*(*t*) is defined by matrix *B*, and the weight matrix between hidden nodes and output is *W*.

The size of our RNN was fixed at 200. We initialized both connection matrix *B* and *J* to be normally distributed (for *B*: *mean* = 0, *SD* = 1/√*N*; for *J*: *mean* = 0, SD = *g*/√*N*; *g* = 1.5), matrix *W* to be all zero, and chose a time constant *τ* = 50 *ms* in the light of previous work ^40, 62^.

All three weights were adjustable and optimized during training. We used the summation of the error function and three regularity terms as a cost function ^44^. The error function was the squared error between the model output and the desired PV. The three regularity terms penalized the magnitude of the averaged firing rate, the intensity of the input and output weights, and the complexity of state trajectories; the hyper-parameters for these three terms in our model were 1e-2, 1, and 1e-2, respectively. The training was finished with PyTorch, and the weights were optimized by Adam (Adaptive Moment Estimation).

To compare the pattern of coefficients, we visualized the three time-varying coefficients as a normalized 3D trajectory, and calculated the Fréchet distance between trajectories of different sessions or monkeys. The distance between monkey C’s array and the RNN was 0.47; in contrast, that between monkey C’s array and single electrode recordings was 0.39, between monkey C’s array and monkey B was 0.56, between monkey C array and monkey G was 0.23. The average distance between monkey C’s array and the permutation was 1.08.

