## Supplementary Figures for "Multiplicative Joint Coding in Preparatory Activity for Reaching Sequence in Macaque Motor Cortex"

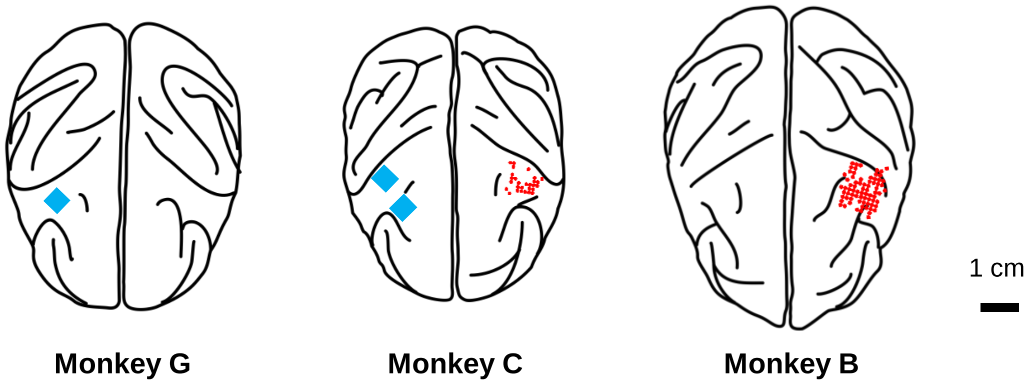


**Supp. Figure 1 Recording locations.**

For monkey G, a 96-channel Utah array was implanted in M1 in the right hemisphere. For monkey C, single-electrode recordings were made from his left hemisphere first. Then two 128-channel Utah arrays were implanted in M1 and PMd of his right hemisphere with a switch of hands. For monkey B, recording sites for single electrodes were located in the left hemisphere. The recorded hemisphere in all monkeys was contralateral to the hand used during the task.

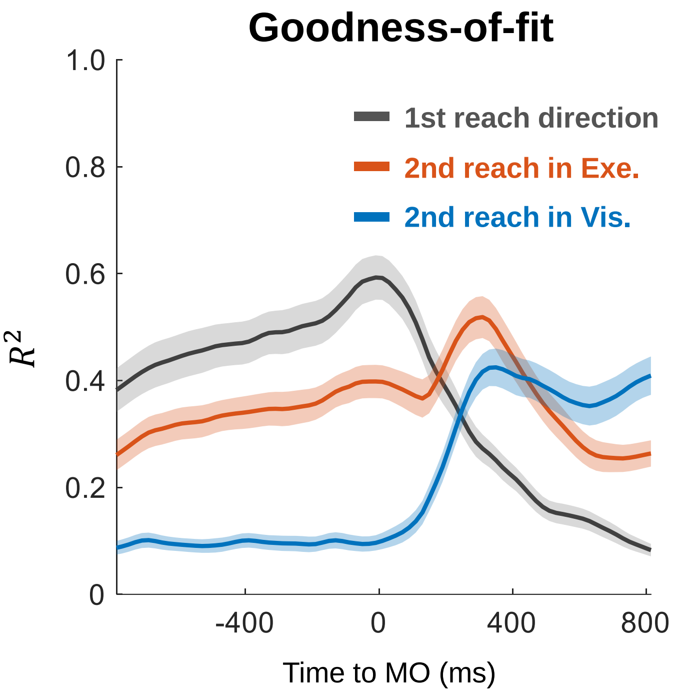


**Supp. Figure 2 Subsequent targets are better fitted in movement reference frame.**

The changing $\boldsymbol{R^2}$ of cosine models in three coordinates was obtained with sliding windows (bin = 200 ms, step = 20 ms): 1st reach direction (gray) fits best before MO; 2nd reach direction in execution coordinates (in Exe., red) fits well throughout the whole trial; 2nd reach direction in visual coordinates (in Vis., blue) fits poorly before MO.

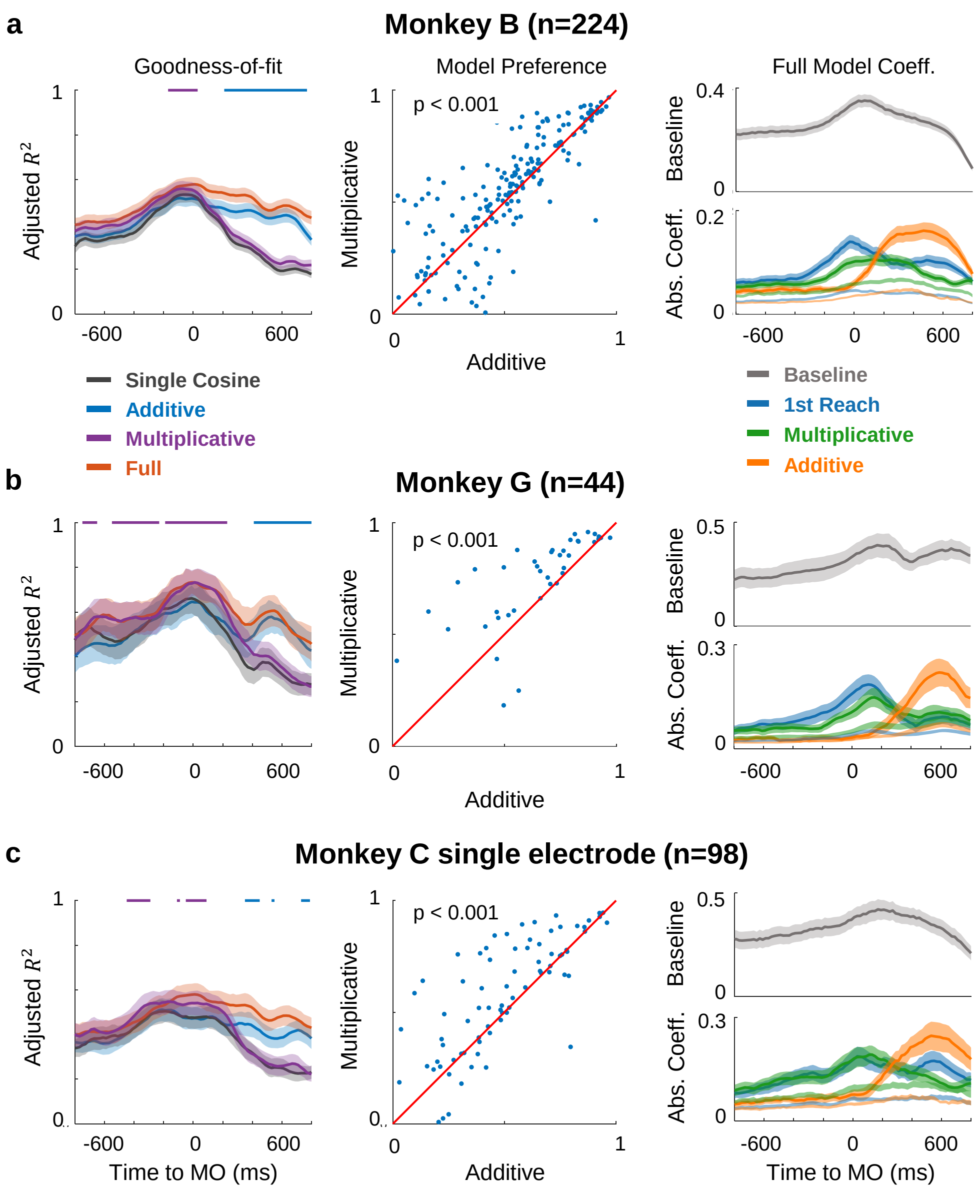


**Supp. Figure 3 Regression performance and coefficient dynamics of other monkeys.**

The same convention as Fig. 5, **a.** Single-electrode recorded data from monkey B. Left: Averaged adjusted $R^2$ for all fitting models in sliding windows. The upper line showed the significance (p<0.0005) of comparison between performance of multiplicative (purple line) and additive (blue line) model. Middle: Scatters of adjusted $R^2$ show that the multiplicative model performed better than the additive model during preparation (-300~-100 ms to MO). Right: Absolute value of each coefficient is averaged across neurons (with twice standard error in shade), the temporal dynamics of which shows the contribution of terms. The coefficient weight of permutation test was plotted in light shade as the chance level. **b.** The results of array data from monkey G. **c.** The results of single electrode recording from monkey C. The time-evolving patterns of model performance are similar among monkeys.

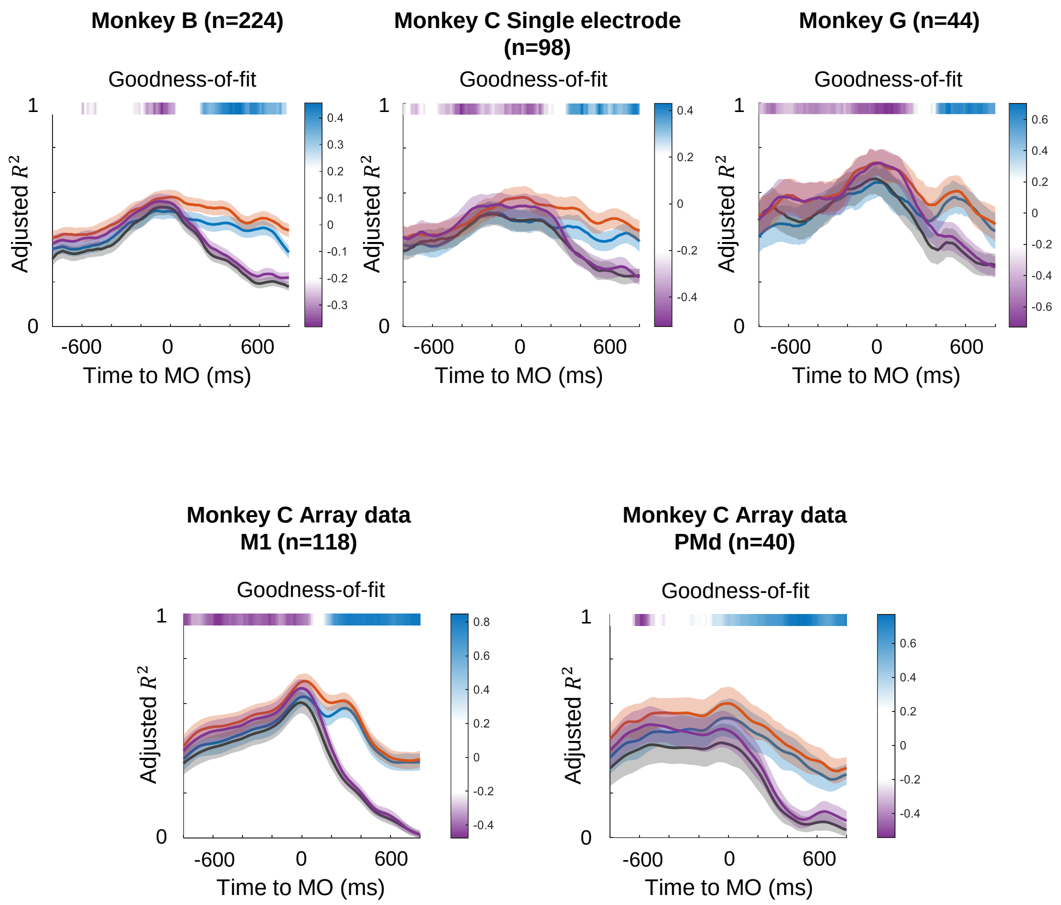


### Supp. Figure 4 Dynamics of goodness-of-fit and effect size

Results of regression are illustrated at the population level for each dataset.: Goodness-of-fit was evaluated with averaged adjusted $R^2$ for all fitting models in a sliding window (with twice standard error in shade). The upper line showed the effect size r (see Methods) of comparison between performance of multiplicative and additive model. When r<0, the line is purple, means multiplicative is superior to additive model; otherwise, blue indicates the additive model is better.

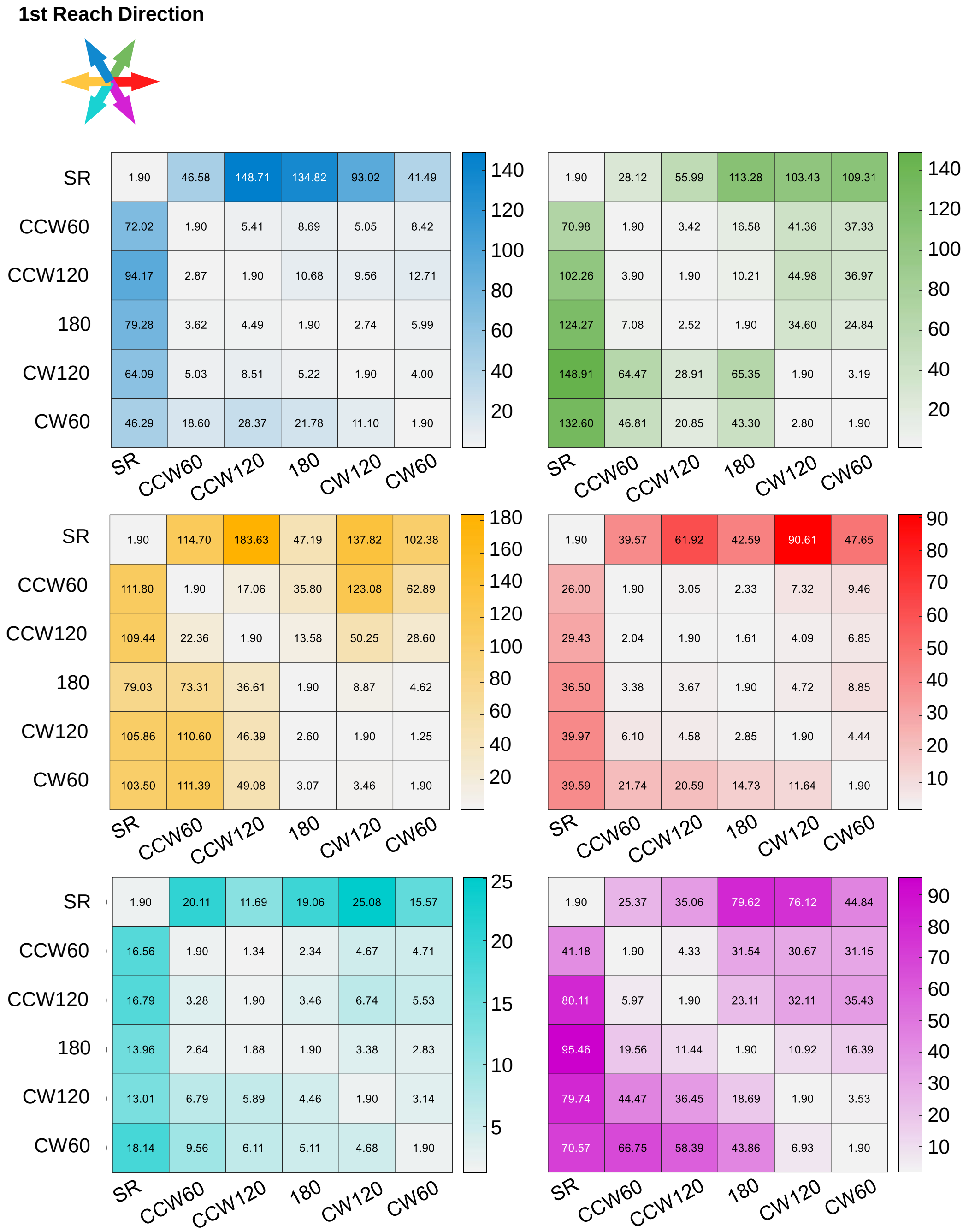


**Supp. Figure 5 Mahalanobis distance of initial state space for each reach direction.**

Mahalanobis distance of the initial state projection in Fig.6c. Rows are the observation clusters and columns are the reference clusters. Color represents the 1st reach direction same as in Fig.6.

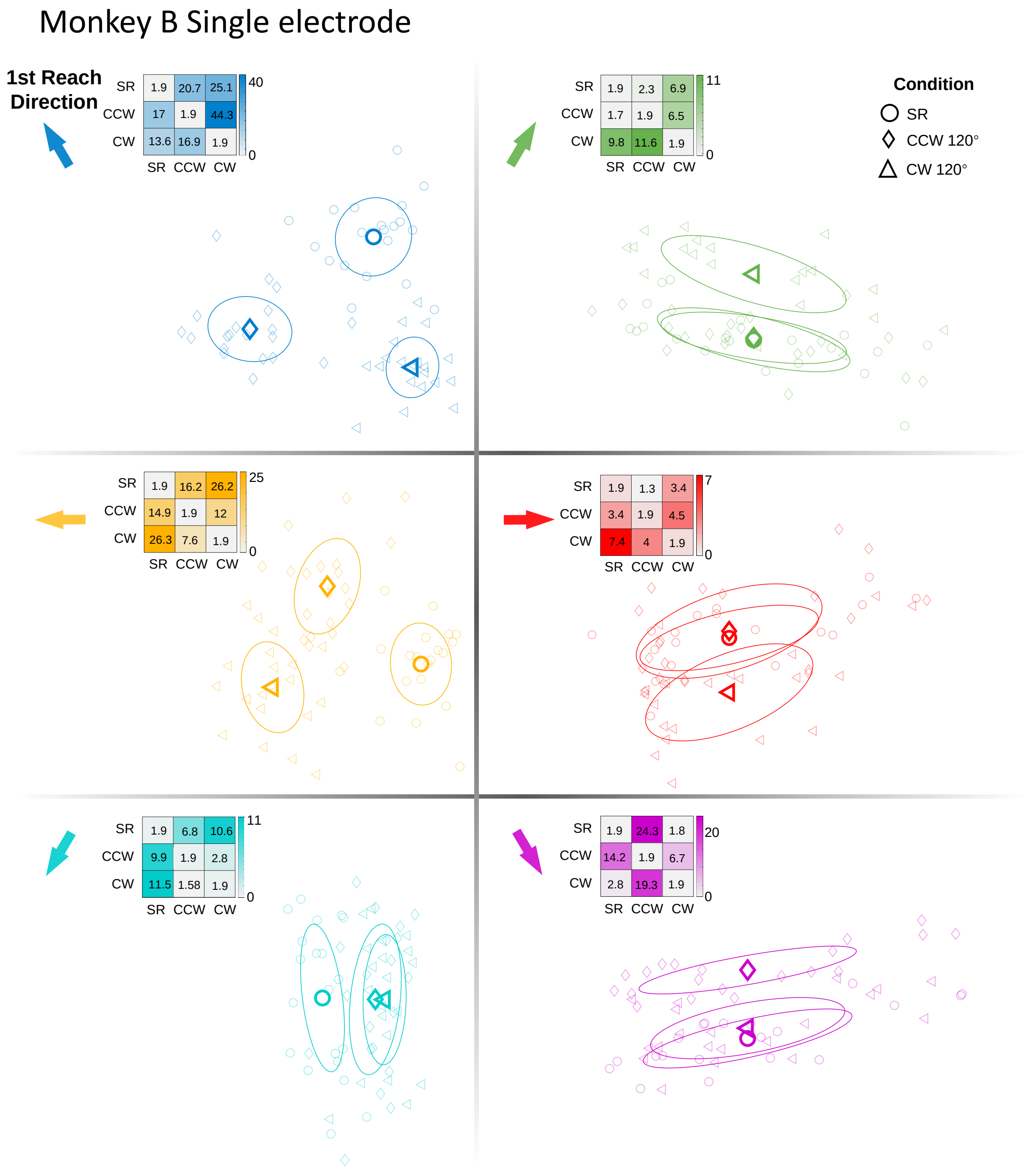


**Supp. Figure 6 Projection of preparatory activity onto PCA-LDA based initial state space and the mahalanobis distance of double reach task data from monkey B single electrode data.**

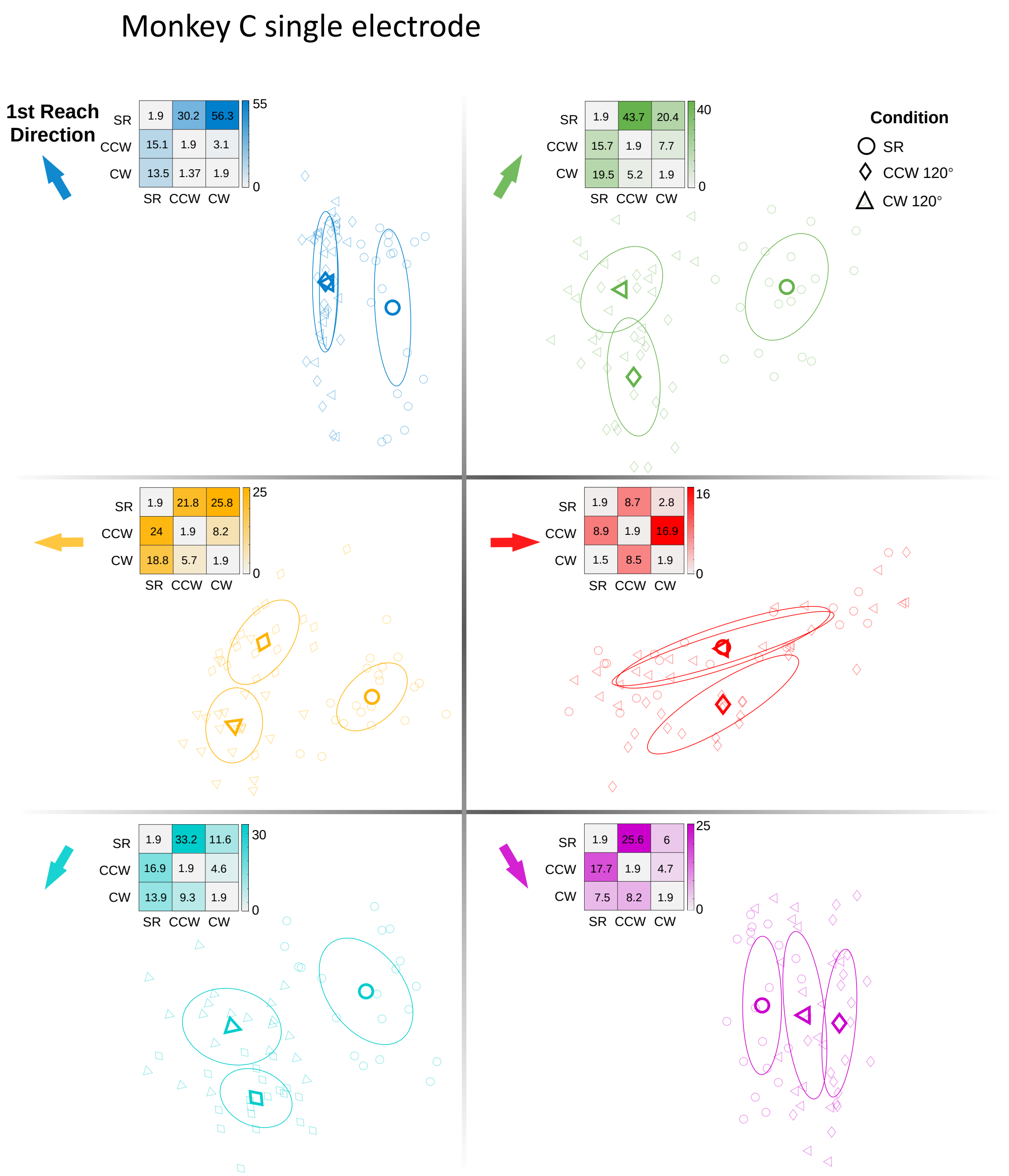


**Supp. Figure 7 Projection of preparatory activity onto PCA-LDA based initial state space and the mahalanobis distance of double reach task data from monkey C single electrode data.**

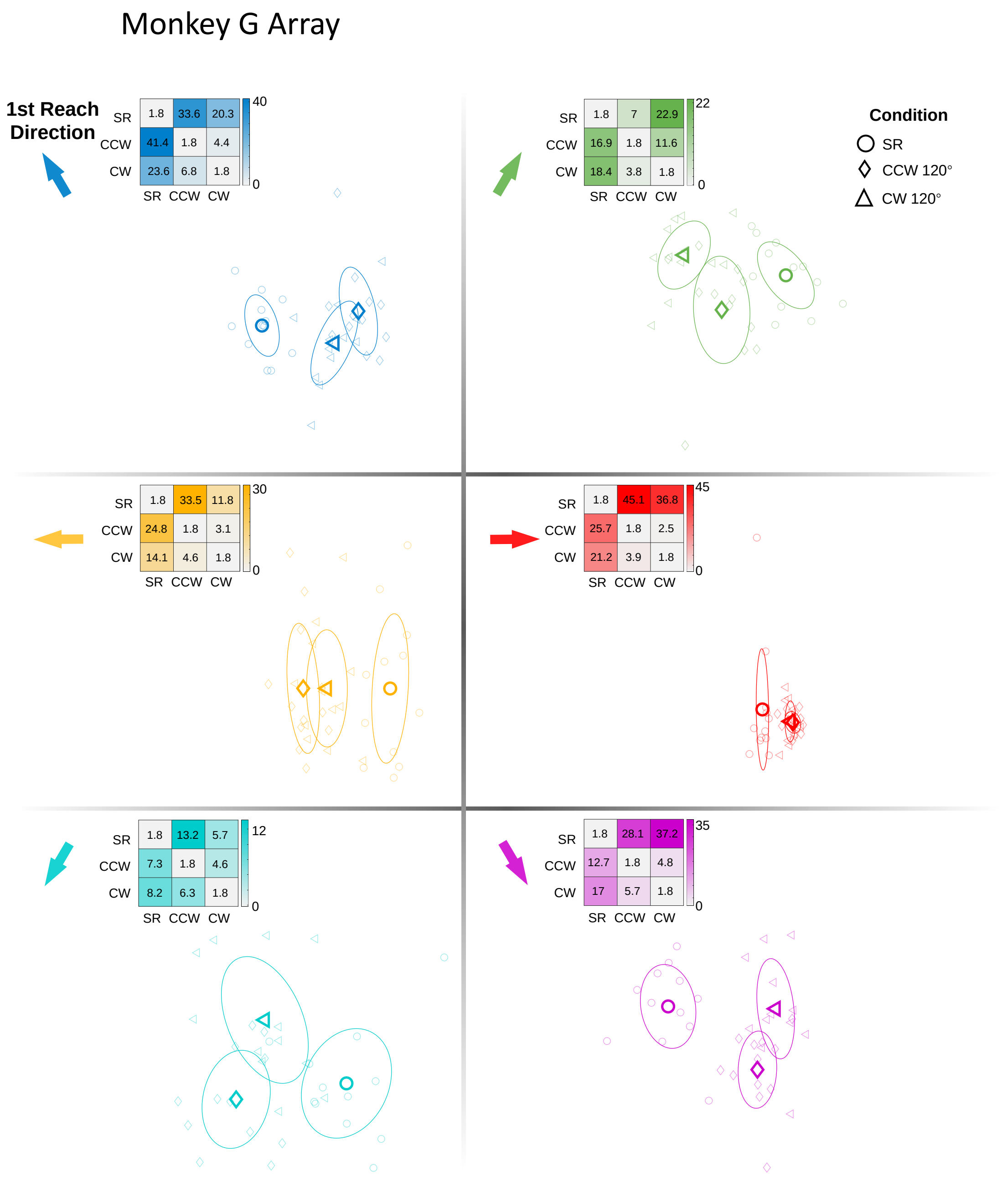


**Supp. Figure 8 Projection of preparatory activity onto PCA-LDA based initial state space and the mahalanobis distance of double reach task data from monkey G array data.**

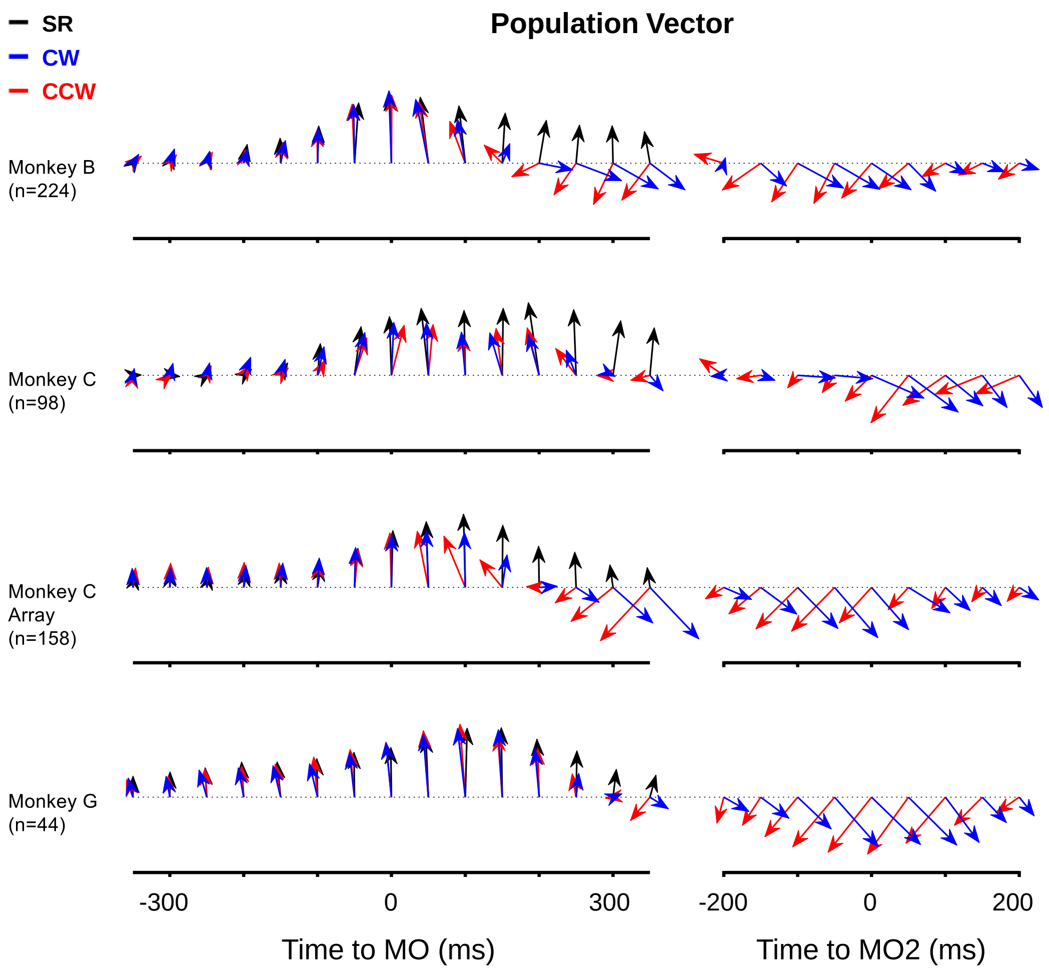


**Supp. Figure 9 Time series of population vectors.**

Time-varying population vectors in SR (black), CW (blue), and CCW (red) conditions. The population vector in DR trials points to the same direction as in SR trials before the 1st reach onset, and then quickly shifts towards the 2nd reaching direction after the 1st movement onset. Therefore, the 1st movement direction can robustly be decoded via linear readout with single gain-modulated neurons.
